# Seeing in the deep: evolution of the opsin gene expression in Bermin crater lake cichlids

**DOI:** 10.1101/2025.02.28.640660

**Authors:** Monika Kłodawska, Adrian Indermaur, Arnold Roger Bitja Nyom, Dmytro Omelchenko, Oldřich Bartoš, Zuzana Musilova

## Abstract

Cichlid visual systems can evolve rapidly during adaptive radiations. This study investigates the Bermin crater lake species flock in Cameroon, comprising thirteen (nine valid and four undescribed) *Coptodon* species, to explore the effects of deep-water light environments on visual evolution. We analyzed visual opsin genes and their expression using 109 retina transcriptomes, focusing on differences among species at varying depths, and seasonal changes in the visual system of a seasonally migrating species. All species exhibit a multichromatic system with at least five cone opsins. While opsin sequence variability among species was minimal due to the flock’s evolutionary youth, opsin expression patterns varied significantly. Deep-water species showed reduced SWS1 and SWS2B expression, consistent with diminished UV-to-violet light in deeper waters. Unexpectedly, we observed increased proportional expression of the red-sensitive LWS opsin gene, contrary to trends seen in other lacustrine fishes. Additionally, in the seasonally deep-dwelling species *Coptodon imbriferna*, opsin expression varied plastically between the rainy (shallow) and dry (deep) seasons, with reduced SWS2B expression when the fish reside in the deeper habitats. To add context of other cichlid systems and to explore shared patterns of molecular adaptation, we compared Bermin cichlids to the deep-water species of the Barombi Mbo crater lake. Both cases exemplify independent evolution of deep-water species, yet their visual systems adapted similarly in the single cones (less UV- and violet-sensitive and more blue-sensitive cones), whereas differently in the long-wavelength sensitive double cones (LWS expression lost in Barombi while increased in Bermin). Overall, our study focuses on an evolutionarily young example of cichlid adaptive radiation, providing a unique opportunity to examine the initial phases of molecular adaptation in visual systems.

## Introduction

### Spectral tunning

Vision is a crucial sense for an organism’s survival. Numerous aquatic species possess specialized adaptations in their visual systems to better suit the light conditions of their habitats (Hauser et al. 2021, Musilova et al. 2021). In fishes, visual spectral sensitivities are often shaped by heterogeneous light environments depending on water clarity and ambient light (Seehausen et al. 2008). Cichlids’ visual system has the evolutionary capacity to rapidly integrate multiple adaptations to the changing light environments (Torres-Dowdall, 2017) or to different habitats, such as deep lacustrine zone (Musilova et al., 2019b). Extensive research of the evolution of East African cichlids’ visual systems revealed multiple mechanisms influencing visual adaptations: mutations in the opsins’ coding sequence, opsin genes expression, and co-expression as well as chromophore usage (Carleton et al. 2016).

The evolution of visual opsin genes has been more dynamic in teleost fishes than in other vertebrate clades, as noticed mainly as the expansion of the visual opsin gene repertoire. On average, teleosts possess six cone opsins genes in their genome, in comparison to two to four in most tetrapods (Cortesi et al. 2015, Musilova et al., 2019a, Lin et al. 2017, Musilova et al. 2021), but in an extreme case of deep-sea spinyfins, there are as many as 40 visual opsins (Musilova et al., 2019a).

Cichlids possess seven cone opsin genes: SWS1 (max. sensitive at 360nm in Nila tilapia), SWS2B (425nm) and SWS2A (456nm), which are expressed in the single cones, and RH2B (472nm), RH2Aβ (518nm), RH2Aα (528nm) and LWS (561nm), which are expressed in the double cones in the retina (Spady et al., 2006). In these two morphological types of photoreceptors, visual pigments are composed of the opsin protein and the chromophore – a light-absorbing component.

The absorption maximum of the light spectrum of the chromophore-opsin complex might be altered by opsin gene mutation leading to the amino acid substitution in the opsin protein (Yokoyama 2008). Some mutations have larger effect on the function, and these so-called key tuning sites tend to be placed in the seven transmembrane regions close to the retinal-binding pocket and usually involve changes in amino acid polarity (Yokoyama and Yokoyama 1990, Chang et al. 1995, Yokoyama and Radlwimmer 1998, Kochendoerfer et al. 1999). Single amino-acid changes might lead to shifts in chromophore-opsin complex sensitivity of between 2 and 35 nm (Asenjo et al.1994; Bowmaker and Hunt 1999; Yokoyama 2000, Carleton and Kocher 2001).

Among all the available visual opsin genes, usually only a subset is expressed in cichlids (Bowmaker 2008, Carleton et al. 2010, Carleton and Yourick 2020, Torres-Dowdall et al. 2021). The overall spectral sensitivities are tuned by different opsins included in these subsets, or by co-expression of multiple opsin genes in individual photoreceptors (Carleton and Kocher 2001; Dalton et al.2014). Hence, their visual systems can vary widely, even among closely related cichlid species within adaptive radiations (Levine et al. 1979, Fernald and Liebman 1980, van der Meer and Bowmaker 1995, Parry et al. 2005, Jordan et al. 2006, Carleton et al. 2016, Musilova et al., 2019b). In Malawi cichlids, for example, the short (expressing SWS1, RH2B, RH2A genes), medium (expressing SWS2B, RH2B, RH2A genes), and long (expressing SWS2A, RH2A, LWS) visual palettes are found in different species albeit having nearly identical opsin genes in their genomes (Carleton et al. 2016, O’Quin et al. 2010). In fact, the long palette with the dominant red-sensitive LWS gene is the most common opsin set found in the ray-finned fishes in general (Lupše et al., 2022). The short and medium palette dominated by green-sensitive opsins have been associated with clear water, nocturnality (Fogg et al. 2022), or deep-water habitat (Musilova et al. 2019b), or has evolved as a result of the LWS gene loss (Lupše et al., 2021; Musilova and Cortesi, 2023), whereas the longer palette is commonly found in species from the turbid, eutrophicated environment, such as Lake Victoria (Seehausen et al. 1997) or rivers.

### Speciation and environment

Cichlids are one of the most diverse teleost families famous for their rapid diversification and adaptation to various ecological niches utilising different depths, foraging modes, food sources, and breeding systems. Adaptive radiation of cichlids, whether in the great African Great lakes such as Malawi (Genner and Turner 2005, Turner 2007), Tanganyika and Victoria (Bezault et al. 2011), in the smaller satellite (Masoko; Talbi et al. 2024) or in crater lakes, such as those in Nicaragua (Torres-Dowdall and Meyer, 2021), or Barombi Mbo (Burress et al. 2017), and Bermin (Stiassny et al., 1992) is often characterized by evolution of highly diversified phenotypes. These adaptations, such as body shape, morphology, or nuptial coloration (Salzburger 2018), have sometimes evolved in enormously short evolutionary timescales. The less diverse and evolutionary younger systems, such as crater lakes in Cameroon with eleven (Barombi Mbo), or nine (plus four undescribed; Bermin) species, or a crater lake Masoko with two ecomorphs of one species provide an excellent opportunity to study visual evolution in greater detail and focusing on the early phases of these adaptive evolutionary processes.

Vision is the primary sensory system that plays an important role in sensing the environment and mediating various behaviors (Hofmann, 2010) and numerous visual adaptations have been associated with different light environments, diet, mate choice, and foraging in cichlids. In relatively young radiation from the Barombi Mbo lake in Cameroon, two deep-water species, convergently shifted their visual sensitivity towards the centre of the light spectrum prevalent in the deep environment (Musilova et al. 2019b). Similarly, Midas cichlids in Nicaraguan lakes, also shifted their visual sensitivity toward shorter wavelengths as compared to the ancestral populations from the turbid environment (Torres-Dowdall, 2017). Speciation through the visual sensory system is well documented in Lake Victoria cichlids, where sexually selected blue and red colour morphs are associated with shallow and deep waters, respectively (Seehausen et al., 2008). In Lake Malawi cichlids a relationship between foraging mode and retinal sensitivity showed that SWS1, UV-sensitive opsin, is significantly associated with foraging preferences. In cichlids foraging on algae and phytoplankton, the overall retinal sensitivity is shifted toward the shorter wavelengths, and they display higher SWS1 expression, than species foraging on fish or invertebrates (Hofmann et al.2009b).

Similar to other crater lakes, Bermin is a deep, relatively oligotrophic lake with species differing partially in the habitats they can be found. While most of the species can be seen in the shallow water, the deeper zones of the lake are inhabitted by a strict deep-water specialist (*Coptodon bythobates*) found only in the deep all year round, by a seasonal species residing in the deep during the dry season but migrating into a shallow waters for spawning in the rainy season (*Coptodon imbriferna*), which strikingly resembles the seasonal species (*Myaka myaka*) in the unrelated endemic radiation in Barombi Mbo. To complete the list, the deep zone of the Bermin lake is also inhabitted by *Coptodon spongotroktis*, which is found commonly in the deep as well as in shallow habitats.

Here, we examine the visual system in thirteen (nine described and four undescribed) species of cichlids from the *Coptodon* genus, from Lake Bermin, Cameroon, West Africa. Bermin is a small crater lake with 700 m in diameter and up to 14 metres deep located in the volcanic chain in the South-West Region of Cameroon. Due to its small size and nine described endemic cichlid species, it is considered to be a place of the highest endemism (Abell et al. 2008). In this study we focus on the molecular mechanisms of the visual adaptation, such as opsin gene diversity and gene expression, of what might be one of the evolutionary youngest examples of adaptive radiation in cichlids, sometimes estimated to be less than ten thousand years old (Stiassny 1992).

## Materials and methods

### Samples collection

Cichlid specimens were collected in the crater lake Bermin, South-West Cameroon, between 2013 and 2017 (research permit numbers: 0000047,49/MINRESI/B00/C00/C10/nye, 0000116,117/MINRESI/B00/C00/C10/C14, 000002-3/MINRESI/B00/C00/C10/C11, 0000032,48 50/MINRESI/B00/C00/C10/C12) during both, dry and rainy seasons. Between 5 and 15 of adult individuals per species were collected each year, covering all 13 species inhabiting Bermin Lake, including four undescribed species. Shallow-water species were captured using gill nets and minnow traps, as well as selective catching with the use of gill nets while snorkeling. Deep-water species, *Coptodon bythobates, C. imbriferna, C. spongotroktis* were captured into gill nets and minnow traps. We also sampled riverine species, *Heterochromis multidens* (SRX6435738), *Hemichromis elongatus* (Batchenga, Sanaga river; SRX6435739), *Etia nguti* (Nguti village; SRX6435784), *Pelmatolapia mariae* (SRX6435737), *Gobiocichla ethelwynnae* (Mamfé city; SRX6435741), *Coptodon kottae* (Barombi Kotto), and *Benitochromis conjunctus* (SRX6435725) in Cameroon, which have been sequenced previously (see provided accession numbers). Fin clips from all the captured specimens have been sampled and stored in 96% EtOH for DNA extraction and further molecular analysis. From freshly euthanized specimens, entire eyeballs were dissected and fixed in RNAlater for the retinal RNA analysis.

### Sanger sequencing, haplotype network construction, and amino-acid diversity

We sequenced 567 single opsins sequences from 109 individuals of Bermin cichlids (partially overlapping with individuals used for the transcriptome sequencing) using a conventional Sanger sequencing method to cover multiple individuals per species for the haplotype network and allelic variability check (Table S2). We complemented our sequence data set by exploring genomes of eleven other Africa cichlids to provide a broader context of opsin evolution (Table 1, Figure 1). DNA was extracted from fin clips stored in 96% EtOH with the Qiagen DNeasy Blood & Tissue kit (www.qiagen.com) following the manufacturer’s protocol. Seven cone opsin genes: SWS1, SWS2B, SWS2A, RH2B, RH2Aβ, RH2Aα, and LWS were Sanger-sequenced for 4-10 individuals per gene per species. The opsin genes were amplified as two (RH2Aα, RH2Aβ), three (SWS1), four (SWS2A), or five (SWS2B, RH2B, LWS) fragments, depending on their length (see Supplementary materials Table S1 for a list of primers used). PCR reactions contained 1 μl of DNA, 5 μl of Master Mix Polymerase (PPP Master Mix, Top-Bio s.r.o.), 0.25 μl of each primer (forward and reverse), 0.25 μl of MgCl and 3.25 μl water up to 10 μl volume. The PCR conditions were initial denaturation: 94°C for 300 s, 35 cycles of denaturation: 94°C for 40 s, annealing between 56°C and 64°C depending on the pair of primers, for 30 s, extension: 72°C for 60 s and after 35 cycles final extension in 72°C for 300 s. PCR products were then purified with ExoSAP-IT PCR Product Cleanup Reagent (Applied Biosystems) and service-sequenced in Macrogen Europe B.V. (https://dna.macrogen.com/eng/). The RH1 genes from Bermin cichlids were obtained from transcriptome data by mapping the transcriptomic libraries to the *C. bakossiorum* RH1 reference gene, allowing extraction of the species-specific RH1 sequences. The obtained sequences were checked for heterozygous sites, trimmed, and aligned to the annotated opsin genes of *Oreochromis niloticus* (GenBank, accession number version MKQE00000000.2) using Geneious 9.1.4 software (https://www.geneious.com, Kearse et al., 2012). Intron sequences were removed for the subsequent analyses. Individual alleles were reconstructed using the PHASE algorithm in DNAsp version 6 (Rozas et al., 2017). To visualize the relationships among the different alleles, haplotype networks have been created for each gene with PopART (Bandelt, Forster, & Röhl, 1999; Leigh & Bryant, 2015) using the minimum spanning network method (ε = 0) and phased sequences from 4 to 10 individuals per species per gene. Additionally, opsin gene sequences were translated into the protein sequence, focusing on amino acid substitutions in the previously identified key spectral tuning amino acid sites (Yokoyama, 2008).

**Figure 1.**
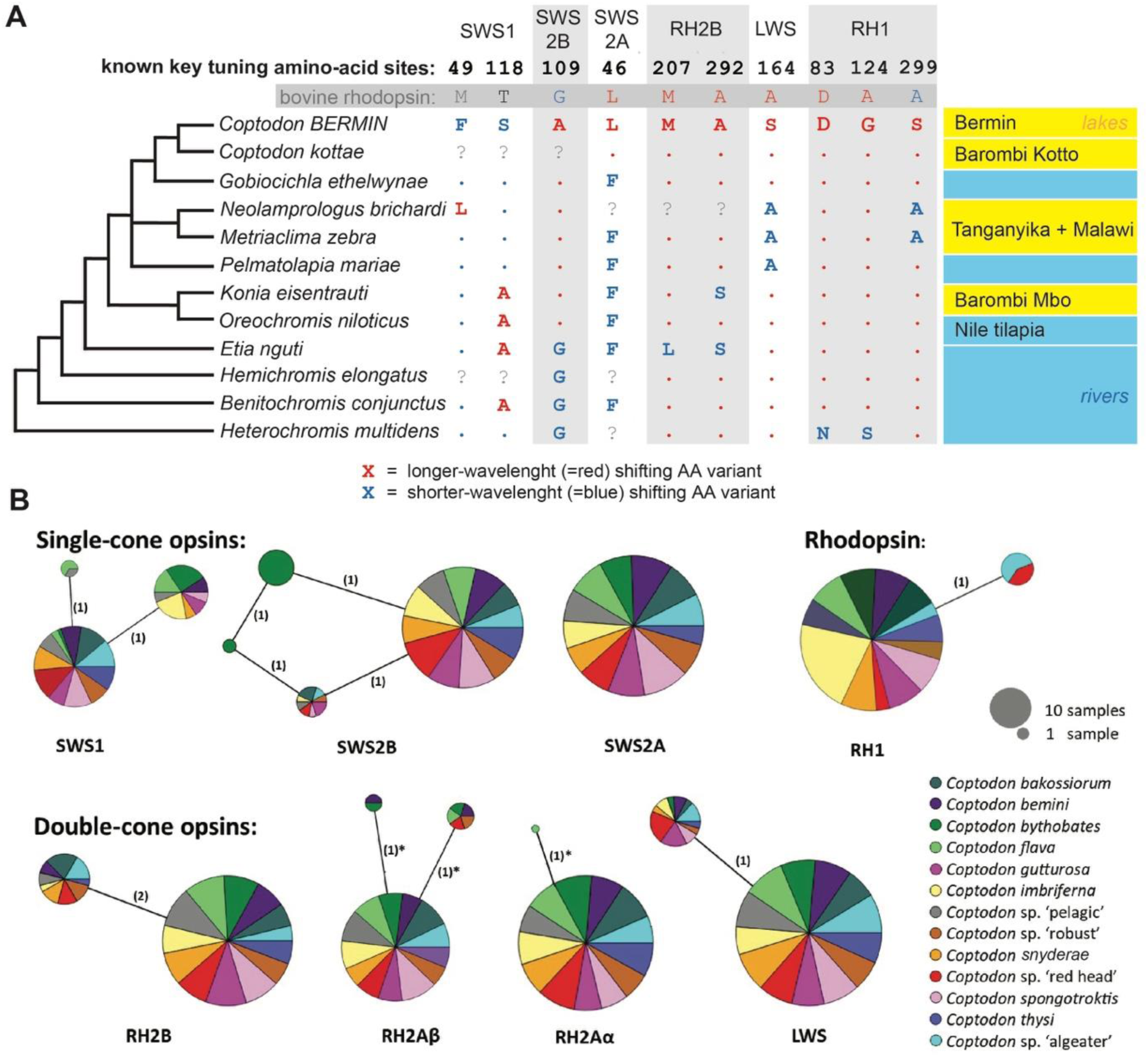
Opsin gene diversity in the context of other African cichlid lineages, and within the Bermin species flock. A) Opsin key tuning amino-acid sites in the cone and rod opsin sequences spanning the African cichlid diversity. Only the variable key tuning sites (as listed in Yokoyama 2008 and Musilova et al., 2019a) are shown. The red and blue colour corresponds to the amino acid variant known to shift the spectral sensitivity (λmax) of the visual pigment towards longer (red), or shorter (blue) wavelengths of the light spectrum. Note that the Bermin cichlids (genus *Coptodon*) possess the red shifted variants in most opsins except for the SWS1 (UV). B) Opsin genes haplotype networks illustrate nucleotide sequence similarity among the 13 Bermin cichlid species. Haplotypes, represented as circles, are sized based on their frequency in the sampled individuals. Mutational steps between haplotypes are indicated by the values next to connecting lines. A star symbol marks the changes which lead to the amino-acid substitution, note that only RH2Aα and RH2Aβ opsins have variable amino acid sites.

**Table 1.** Samples used for transcriptome sequencing.

| DNA code | Species | Sex | Season | Date of collection | Accession number | note | source |
| --- | --- | --- | --- | --- | --- | --- | --- |
| 41D 1 | *Coptodon* sp. 'algaeater' | M | dry | December 2014 | tba |  | this study |
| 58E2 | *Coptodon* sp. 'algaeater' | M | dry | February 2017 | tba |  | this study |
| 58E3 | *Coptodon* sp. 'algaeater' | F | dry | February 2017 | tba |  | this study |
| 41C9 | *Coptodon* sp. 'algaeater' | M | dry | December 2014 | tba |  | this study |
| 41D 2 | *Coptodon* sp. 'algaeater' | F | dry | December 2014 | tba |  | this study |
| 40I6 | *Coptodon* sp. 'algaeater' | M | dry | December 2014 | tba |  | this study |
| 40A 1 | *Coptodon bakossiorum* | F | dry | December 2014 | tba | DGE | this study |
| 40A 3 | *Coptodon bakossiorum* | M | dry | December 2014 | tba |  | this study |
| 40A 2 | *Coptodon bakossiorum* | M | dry | December 2014 | tba |  | this study |
| 41A 4 | *Coptodon bakossiorum* | F | dry | December 2014 | tba | excluded (low quality) | this study |
| 58G 1 | *Coptodon bakossiorum* | M | dry | February 2017 | tba | DGE | this study |
| 58G 4 | *Coptodon bakossiorum* | M | dry | February 2017 | tba | DGE | this study |
| 58G 3 | *Coptodon bakossiorum* | F | dry | February 2017 | tba |  | this study |
| 58B3 | *Coptodon bemini* | M | dry | February 2017 | SAMN29579 949 |  | Lupše et al. 2022 |
| 40E2 | *Coptodon bemini* | F | dry | December 2014 | SAMN29164 965 |  | Lupše et al. 2022 |
| 40E1<br>— | *Coptodon*<br>*bemini* | M | dry | December<br>2014 | SAMN30522<br>615 |  | Lupše et al.<br>2022 |
| 83I2 | *Coptodon*<br>*bemini* | M | rainy | July 2017 | SAMN30336<br>809 |  | Lupše et al.<br>2022 |
| 40E3<br>— | *Coptodon*<br>*bemini* | M | dry | December<br>2014 | SAMN29176<br>028 |  | Lupše et al.<br>2022 |
| 40B9 | *Coptodon*<br>*bemini* | M | dry | December<br>2014 | SAMN29162<br>646 |  | Lupše et al.<br>2022 |
| 83D<br>1 | *Coptodon*<br>*bemini* | M | rainy | July 2017 | SAMN29164<br>340 |  | Lupše et al.<br>2022 |
| 83I1 | *Coptodon*<br>*bemini* | M | rainy | July 2017 | tba |  | this study |
| 63F8 | *Coptodon*<br>*bythobates* | no<br>info | dry | March<br>2017 | tba |  | this study |
| 51A<br>3 | *Coptodon*<br>*bythobates* | M | dry | March<br>2016 | tba |  | this study |
| 51A<br>8 | *Coptodon*<br>*bythobates* | F | dry | March<br>2016 | tba |  | this study |
| 51A<br>5 | *Coptodon*<br>*bythobates* | M | dry | March<br>2016 | tba | DGE | this study |
| 83D<br>7 | *Coptodon*<br>*bythobates* | M | rainy | July 2017 | tba |  | this study |
| 83F2 | *Coptodon*<br>*bythobates* | M | rainy | July 2017 | tba |  | this study |
| 58F5 | *Coptodon*<br>*bythobates* | M | dry | February<br>2017 | tba | DGE | this study |
| 58F4 | *Coptodon*<br>*bythobates* | M | dry | February<br>2017 | tba | DGE | this study |
| 58D<br>1 | *Coptodon*<br>*flava* | M | dry | February<br>2017 | SAMN31148<br>890 |  | Lupše et al.<br>2022 |
| 58D<br>2 | *Coptodon*<br>*flava* | M | dry | February<br>2017 | SAMN31149<br>648 |  | Lupše et al.<br>2022 |
| 58D<br>3 | *Coptodon*<br>*flava* | F | dry | February<br>2017 | SAMN31149<br>921 |  | Lupše et al.<br>2022 |
| 83B6 | *Coptodon*<br>*flava* | no<br>info | rainy | July 2017 | SAMN31156<br>807 |  | Lupše et al.<br>2022 |
| 83D<br>4 | *Coptodon*<br>*flava* | M | rainy | July 2017 | tba |  | this study |
| 83A<br>6 | *Coptodon*<br>*flava* | no<br>info | rainy | July 2017 | SAMN31156<br>302 |  | Lupše et al.<br>2022 |
| 40D<br>4 | *Coptodon*<br>*flava* | M | dry | December<br>2014 | SAMN31155<br>141 |  | Lupše et al.<br>2022 |
| 41B7 | *Coptodon*<br>*flava* | M | dry | December<br>2014 | SAMN31157<br>679 |  | Lupše et al.<br>2022 |
| 83D<br>5 | *Coptodon*<br>*flava* | F | rainy | July 2017 | SAMN31157<br>433 |  | Lupše et al.<br>2022 |
| 41B8 | *Coptodon*<br>*flava* | M | dry | December<br>2014 | SAMN31148<br>813 |  | Lupše et al.<br>2022 |
| 40C3 | *Coptodon*<br>*gutturosa* | M | dry | December<br>2014 | tba |  | this study |
| 58C1 | *Coptodon gutturosa* | M | dry | February 2017 | tba |  | this study |
| 40E6 | *Coptodon gutturosa* | F | dry | December 2014 | tba |  | this study |
| 58C3 | *Coptodon gutturosa* | F | dry | February 2017 | tba | DGE | this study |
| 58C2 | *Coptodon gutturosa* | M | dry | February 2017 | tba |  | this study |
| 63G8 | *Coptodon gutturosa* | M | dry | March 2017 | tba | DGE | this study |
| 40F2 | *Coptodon gutturosa* | F | dry | December 2014 | tba |  | this study |
| 40E9 | *Coptodon gutturosa* | M | dry | December 2014 | tba | DGE | this study |
| _83E5 | *Coptodon imbriferna* | M | rainy | July 2017 | SAMN30548120 | DGEII | Lupše et al. 2022 |
| _83E6 | *Coptodon imbriferna* | M | rainy | July 2017 | SAMN30594652 | DGEII | Lupše et al. 2022 |
| .83E7_ | *Coptodon imbriferna* | F | rainy | July 2017 | tba | DGEII | this study |
| .83E9_ | *Coptodon imbriferna* | F | rainy | July 2017 | tba | DGEII | this study |
| 41C7 | *Coptodon imbriferna* | F | dry | December 2014 | tba | DGEII | this study |
| 41C8 | *Coptodon imbriferna* | M | dry | December 2014 | SAMN30524742 |  | Lupše et al. 2022 |
| 44D7 | *Coptodon imbriferna* | F | dry | March 2016 | tba | DGEII | this study |
| 44E8_ | *Coptodon imbriferna* | M | dry | March 2016 | tba | DGEII | this study |
| 44E9_ | *Coptodon imbriferna* | M | dry | March 2016 | tba |  | this study |
| 44F3 | *Coptodon imbriferna* | F | dry | March 2016 | tba | DGEII | this study |
| 44F4 | *Coptodon imbriferna* | F | dry | March 2016 | tba |  | this study |
| 58A1 | *Coptodon imbriferna* | M | dry | February 2017 | SAMN31146633 | DGE, DGEII | Lupše et al. 2022 |
| 58A6 | *Coptodon imbriferna* | M | dry | February 2017 | SAMN31184165 | DGEII | Lupše et al. 2022 |
| 58F7 | *Coptodon imbriferna* | M | dry | February 2017 | SAMN30540913 | excluded (low quality) | Lupše et al. 2022 |
| 58F8 | *Coptodon imbriferna* | F | dry | February 2017 | SAMN31146607 | excluded (low quality) | Lupše et al. 2022 |
| 58F9 | *Coptodon imbriferna* | M | dry | February 2017 | SAMN30541242 | DGE | Lupše et al. 2022 |
| 83A1 | *Coptodon imbriferna* | no info | rainy | July 2017 | SAMN30541253 |  | Lupše et al. 2022 |
| 83A3 | *Coptodon imbriferna* | no info | rainy | July 2017 | tba |  | this study |
| 83A<br>4 | *Coptodon*<br>*imbriferna* | no<br>info | rainy | July 2017 | tba |  | this study |
| 83C8 | *Coptodon*<br>*imbriferna* | F | rainy | July 2017 | SAMN30547<br>393 | DGEII | Lupše et al.<br>2022 |
| 83C9 | *Coptodon*<br>*imbriferna* | M | rainy | July 2017 | SAMN30547<br>878 |  | Lupše et al.<br>2022 |
| 58H<br>5 | *Coptodon*<br>*sp. 'pelagic'* | F | dry | February<br>2017 | tba |  | this study |
| 58H<br>6 | *Coptodon*<br>*sp. 'pelagic'* | F | dry | February<br>2017 | tba |  | this study |
| 63G<br>6 | *Coptodon*<br>*sp. 'pelagic'* | M | dry | March<br>2017 | tba |  | this study |
| 63G<br>7 | *Coptodon*<br>*sp. 'pelagic'* | M | dry | March<br>2017 | tba | excluded (low<br>quality) | this study |
| 40I4 | *Coptodon*<br>*sp. 'pelagic'* | M | dry | December<br>2014 | tba |  | this study |
| 40H<br>7 | *Coptodon*<br>*sp. 'pelagic'* | M | dry | December<br>2014 | tba |  | this study |
| 51C3 | *Coptodon*<br>*sp. 'pelagic'* | F | dry | March<br>2016 | tba |  | this study |
| 51D<br>8 | *Coptodon*<br>*sp.red head* | M | dry | March<br>2016 | tba |  | this study |
| 58B6 | *Coptodon*<br>*sp.red head* | M | dry | February<br>2017 | tba |  | this study |
| 58B7 | *Coptodon*<br>*sp.red head* | M | dry | February<br>2017 | tba |  | this study |
| 58B9 | *Coptodon*<br>*sp.red head* | M | dry | February<br>2017 | tba |  | this study |
| 58B8 | *Coptodon*<br>*sp.red head* | M | dry | February<br>2017 | tba |  | this study |
| 51D<br>7 | *Coptodon*<br>*sp.red head* | M | dry | March<br>2016 | tba |  | this study |
| 63G<br>4 | *Coptodon*<br>*sp. 'robust'* | M | dry | March<br>2017 | tba |  | this study |
| 63G<br>5 | *Coptodon*<br>*sp. 'robust'* | M | dry | March<br>2017 | tba |  | this study |
| 41E7<br>_ | *Coptodon*<br>*sp. 'robust'* | M | dry | December<br>2014 | tba |  | this study |
| 51C7 | *Coptodon*<br>*sp. 'robust'* | no<br>info | dry | March<br>2016 | tba |  | this study |
| 51C8 | *Coptodon*<br>*sp. 'robust'* | no<br>info | dry | March<br>2016 | tba |  | this study |
| 63D<br>9 | *Coptodon*<br>*snyderae* | F | dry | March<br>2017 | SAMN31167<br>505 |  | Lupše et al.<br>2022 |
| 63E1<br>_ | *Coptodon*<br>*snyderae* | M | dry | March<br>2017 | SAMN31170<br>753 |  | Lupše et al.<br>2022 |
| 44H<br>2 | *Coptodon*<br>*snyderae* | M | dry | March<br>2016 | SAMN31164<br>897 |  | Lupše et al.<br>2022 |
| 63E5 | *Coptodon*<br>*snyderae* | M | dry | March<br>2017 | SAMN31170<br>756 |  | Lupše et al.<br>2022 |
| 58I7 | *Coptodon*<br>*snyderae* | no<br>info | dry | February<br>2017 | SAMN31170<br>755 |  | Lupše et al.<br>2022 |
| 51B6 | *Coptodon*<br>*snyderae* | M | dry | March<br>2016 | SAMN31167<br>504 |  | Lupše et al.<br>2022 |
| 51B5 | *Coptodon*<br>*snyderae* | M | dry | March<br>2016 | SAMN31164<br>898 |  | Lupše et al.<br>2022 |
| 51B7 | *Coptodon*<br>*snyderae* | F | dry | March<br>2016 | SAMN31170<br>754 |  | Lupše et al.<br>2022 |
| 41E8<br>– | *Coptodon*<br>*spongotrok*<br>*tis* | M | dry | December<br>2014 | tba |  | this study |
| 63F9 | *Coptodon*<br>*spongotrok*<br>*tis* | F | dry | March<br>2017 | tba |  | this study |
| 58B2 | *Coptodon*<br>*spongotrok*<br>*tis* | M | dry | February<br>2017 | tba |  | this study |
| 63H<br>8 | *Coptodon*<br>*spongotrok*<br>*tis* | M | dry | March<br>2017 | tba |  | this study |
| 44H<br>9 | *Coptodon*<br>*spongotrok*<br>*tis* | M | dry | March<br>2016 | tba |  | this study |
| 63H<br>9 | *Coptodon*<br>*spongotrok*<br>*tis* | F | dry | March<br>2017 | tba |  | this study |
| 58B1 | *Coptodon*<br>*spongotrok*<br>*tis* | M | dry | February<br>2017 | tba |  | this study |
| 44I1 | *Coptodon*<br>*spongotrok*<br>*tis* | F | dry | March<br>2016 | tba |  | this study |
| 63D<br>6 | *Coptodon*<br>*thysi* | M | dry | March<br>2017 | SAMN31181<br>402 |  | Lupše et al.<br>2022 |
| 63D<br>7 | *Coptodon*<br>*thysi* | M | dry | March<br>2017 | SAMN31181<br>403 |  | Lupše et al.<br>2022 |
| 63D<br>8 | *Coptodon*<br>*thysi* | F | dry | March<br>2017 | SAMN31181<br>404 |  | Lupše et al.<br>2022 |
| 41E3<br>– | *Coptodon*<br>*thysi* | M | dry | December<br>2014 | SAMN31172<br>241 |  | Lupše et al.<br>2022 |
| 63H<br>7 | *Coptodon*<br>*thysi* | M | dry | March<br>2017 | SAMN31183<br>056 |  | Lupše et al.<br>2022 |
| 63D<br>5 | *Coptodon*<br>*thysi* | F | dry | March<br>2017 | SAMN31183<br>055 |  | Lupše et al.<br>2022 |
| –41E<br>4 | *Coptodon*<br>*thysi* | F | dry | December<br>2014 | SAMN31183<br>054 |  | Lupše et al.<br>2022 |

### Transcriptome sequencing

We sequenced 109 individual retina transcriptomes of all Bermin species (Table 1). The retinas were dissected from the eyeballs preserved in the RNAlater solution. RNA was extracted from the dissected retina using Qiagen RNeasy Micro Kit (www.qiagen.com). The extracted RNA concentration and integrity were verified using a 2100 Bioanalyzer (Agilent) and Qubit Fluorometer (Thermofisher Scientific). Libraries were prepared in-house with NEBNext® Ultra™ RNA Library Prep Kit for Illumina® (New England BioLabs Inc.) and service-sequenced with Novogene Co., Ltd. Prepared libraries covered 5 – 11 individuals per species. Additional information regarding the specimens’ sex and seasons during which they have been collected are listed in Table T1. Sequenced libraries were quality-checked and trimmed using FastQC (Andrews 2010) and Trim Galore ver.0.4.4 (https://www.bioinformatics.babraham.ac.uk/projects/trim_galore/) respectively.

### Opsin gene expression analysis and statistical tests

Relative opsin gene expression was calculated using Geneious software 9.1.4. For each sample, the pair-end library has been mapped against the reference dataset made of 7 cone opsin genes (SWS1, SWS2B, SWS2A, RH2B, RH2Aβ, RH2Aα, LWS) from one selected species, *Coptodon bakossiorum*. Additionally, to distinguish between similar RH2Aβ and RH2Aα genes, we subsequently mapped each library against the 4^th^ exon of RH2Aβ and RH2Aα genes (where we found 10 SNPs that are different between RH2Aβ and RH2Aα in Bermin cichlids) to get the ratio between these two genes. Based on the number of mapped reads to each reference gene, we calculated the FPKM (fragments per kilobase and million reads) taking into account also the library size and the length of each gene. We then calculated the proportional expression of opsin genes in percentage of all opsins. These calculations of proportional opsin gene expression were performed separately within the single and the double cone opsins. Additionally, to check whether Bermin cichlids utilize A1 retinal or a mixture of A1 and A2 retinals which would shift the sensitivity of an opsin protein and retinal complex sensitivity to the longer wavelength and help to accurately estimate λmax, we mapped all the libraries against Cyp27c1 *Coptodon bakossiorum* reference (obtained after mapping the reads against the Cyp27c1 gene from *Danio rerio*; acc. no: NM_001113337). To calculate the potential expression level of Cyp27c1 we also mapped the libraries to several housekeeping genes: ube2z, ef2z, actb (from *Coptodon bakossiorum*–) and gnat2 (from *Oreochromis niloticus*).

We used beta regression analysis to formally test for significant differences among the proportional expression between deep-dwelling species (*C. bythobates, C. imbriferna, C. spongotroktis*) and shallow-water species. The same method was also used for the comparison of the rainy season and dry season samples in *Coptodon imbriferna*. We used the R package betareg (Cribari-Neto & Zeileis 2010), which allows handling of non-transformed data. This analysis is suitable to fit the dependent variable (in our case the proportional expression of each opsin gene) in the interval (0,1). We tested the effect of depth for selected cone opsin gene class separately (i.e, SWS1, SWS2B, SWS2A, RH2Ab, and LWS based on the selection and hypotheses formulated based on the order species by expression as seen in Figure 2B and Supplementary TableS5). We tested the effect of season and sex in the seasonal migratory species, *C. imbriferna*, on the expression of SWS1, SWS2B, RH2Ab, RH2Aa, and LWS (see Figure 4 for visualized results). In this case, SWS2A was basically showing a mirror pattern to SWS2B (since it’s a complementary gene in single cones) and was not tested separately.

**Figure 2.**
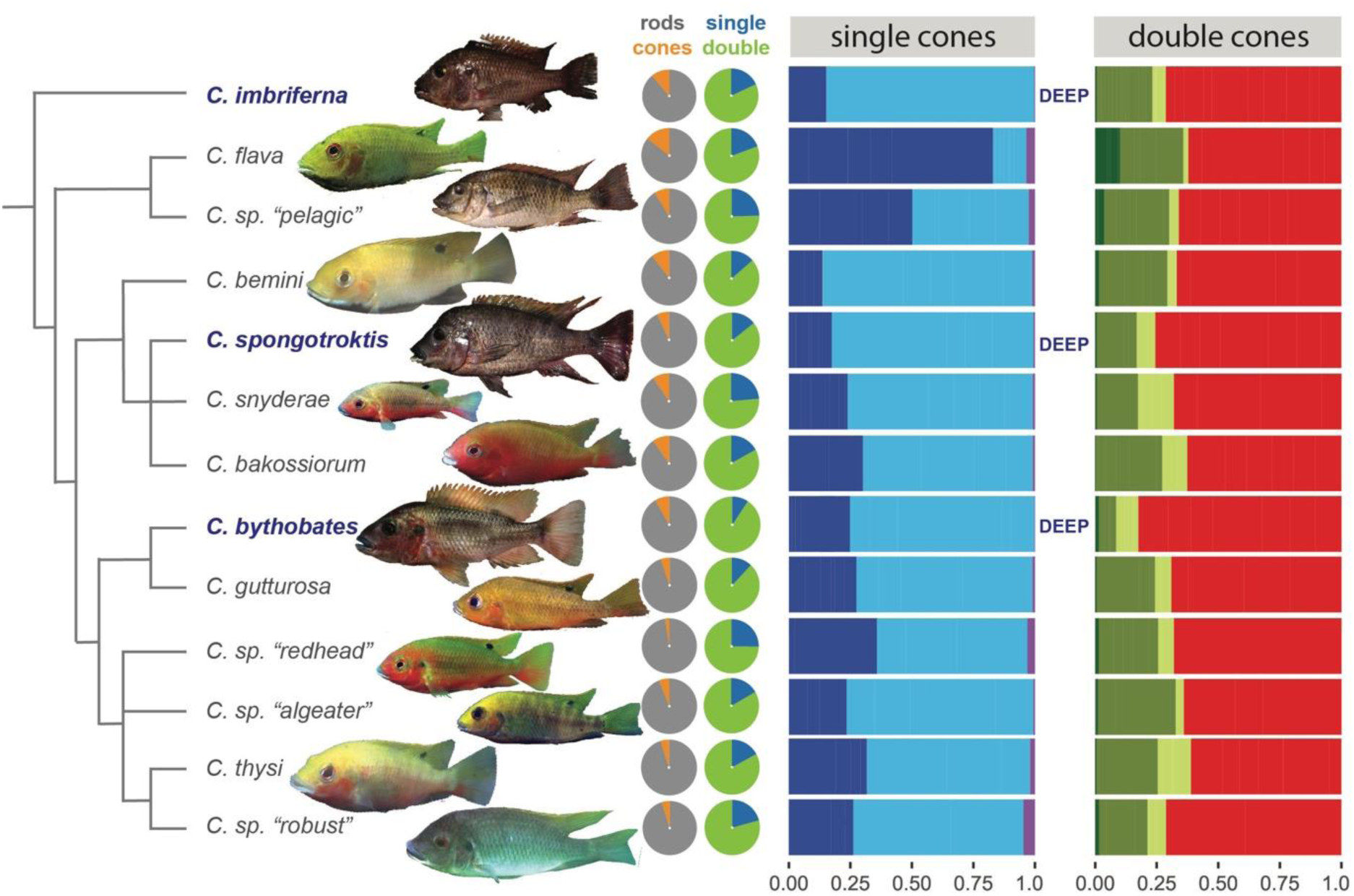
Opsin gene expression in Bermin cichlids. A proportional cone opsin gene expression profiles of the retina of all 13 cichlid species inhabiting the Bermin crater lake based on the retinal transcriptome analysis. The species average values are shown as proportions of the total single and double cone opsin expression (separately). For the values per individual see Table 2, Supplementary Table S4 and S5, and the Supplementary Figure S1.

To somewhat present the overall retina sensitivity, we show single and double cone sensitivity averages, as per Hofman et al.2009b. We calculated the weighted average of all seven opsin genes expression within single and double cones separately, following the equation in Hofman et al.2009b, and we used previously, experimentally measured values of cone opsin genes sensitivities in tilapia, λmax, i.e. SWS1 (360nm), SWS2B (425nm), SWS2A (456nm), RH2B (472nm), RH2Aβ (518nm), RH2Aα (528nm) and LWS (561nm) (Spady et al., 2006). In a similar fashion we compared average single and double cone sensitivity between Bermin and Barombi Mbo cichlids as both flocks are sympatric and share similarities such as comparable species counts and inhabiting crater lakes in close proximity, albeit with differing ages; Barombi Mbo cichlids are relatively older (hundred thousand to 1 million years), compared to ten(s) thousand(s) years of age of Bermin cichlids.

### Differential gene expression

To investigate differential gene expression, we applied a comparative transcriptomics pipeline across two distinct analyses: lake Bermin deep vs. shallow species comparison, and seasonality (and sex) comparison in *C. imbriferna.* In the former case, we aimed to identify genes differentially expressed between deep-water and shallow-water Bermin species, following a comparative transcriptomics approach similar to Musilova et al. (2019b). Here we compared retinal transcriptomes from adult individuals from two shallow-water species (*C. bakossiorum* and *C. gutturosa*) and two deep-water species (*C. bythobates* and *C. imbriferna*). In the latter analysis aiming to assess seasonal effects and detach it from the sex-specific expression, we analyzed gene expression across four groups of *C. imbriferna*: males and females collected during the dry and rainy seasons. This analysis used the same transcriptomics pipeline to examine potential seasonal and sex-related gene expression differences. The transcriptomes selected and used in the DGE analysis are marked in Table 1.

The quality of sequenced libraries was assessed using FastQC (Andrews, 2010), and reads were trimmed to remove low-quality bases and adapter sequences using Trim Galore v0.4.4 (https://www.bioinformatics.babraham.ac.uk/projects/trim_galore/). Reads were additionally filtered for PhiX contamination and residual adapter sequences using bbduk.sh. Filtered reads were aligned to the Nile tilapia reference genome using BWA, with no read deduplication performed. Gene-level coverage was calculated using bedtools multicov. DESeq2 was used to identify differentially expressed genes (DEGs) across species comparisons, setting thresholds for statistical significance and log2 fold change. Venn diagrams were created for deep-water and shallow-water comparisons to visualize DEGs. After DESeq2 analysis, genes were filtered for statistical significance (adjusted p-value < 0.1) and directional log2 fold change (negative for downregulated, positive for upregulated). DEG lists from each species comparison were processed, and the intersections were visualized with ggvenn, using distinct colors for each comparison and setting a false discovery rate of 0.05 and logFC threshold of 1.

## Results and discussion

In this study, we investigated molecular mechanisms of the visual system adaptation in cichlids from the Bermin crater lake in Cameroon. Thanks to the relatively small size of this species flock, we were able to cover all 13 species (nine described plus four undescribed) from the Bermin radiation, and examine its visual system in greater detail. We analyzed both DNA and RNA to provide a comprehensive overview of visual evolution in the Bermin species flock. We believe that comparable adaptive radiations—such as those in Barombi Mbo, Masoko, and the Nicaraguan crater lakes—will eventually allow for cross-system comparisons, identifying general mechanisms of adaptation at the initial stages of diversification.

Our results, based on 109 sequenced retina transcriptomes and over 500 individual opsin gene sequences, show that visual system adaptation occurs primarily at the level of gene expression. We found differences in relative opsin gene expression that correlate with the depth at which different species occur. Additionally, we report seasonal changes in expression between the dry and rainy seasons in a seasonal spawner, *C. imbriferna*. In contrast, there is virtually no variation in opsin nucleotide sequences, as evidenced by over 100 individuals sequenced for all seven cone opsin genes (Supplementary Table S2).

### Nucleotide diversity in the opsin gene sequences

Bermin Coptodons, like other African cichlids, possess seven cone opsin genes in their genome (Carleton et al. 2016). We sequenced all seven cone opsin genes to explore nucleotide diversity within the entire Bermin flock. We found very low or no variation in opsin gene haplotypes, with SWS1, SWS2B, SWS2A, RH2B, and LWS also showing no variability in amino acid sequences (Figure 1B). Within all species, only the two green-sensitive RH2Aα and RH2Aβ genes exhibited multiple alleles differing in their protein sequences (Figure 1B). None of the variable amino acids (i.e., 10 and 159; position respective to the bovine rhodopsin) are located in known key tuning sites or any transmembrane region, and thus, we do not expect major functional changes to result from these substitutions.

In the evolutionary older fish assemblages, such as Tanganyika cichlids (Ricci et al., 2022) or Baikal sculpins (Hunt et al., 1996), amino acid mutations also alter the sensitivity of the rhodopsin genes in the deep-water species. This is not the case in the much younger Bermin species flock with only one amino acid sequence across all species for most opsins (Figure 1B).

The low sequence variation aligns with findings from other crater lake systems, such as the Barombi Mbo radiation (2–5 amino acid variants per gene; Musilova et al., 2019b) and Midas cichlids from Nicaragua (Torres-Dowdall et al. 2017). The Bermin species flock exhibits the lowest diversity among these systems, somewhat comparable to the Pundamilia genus from Lake Victoria (0–4 variable amino acid sites; Carleton et al. 2005). This is most likely due to its young evolutionary age, estimated to be as little as eight thousand years (Stiassny, 1992), although the crater lake itself is estimated to be much less than a million years old (Thieme et al., 2005).

A precise estimate of the species flock’s age is still lacking, and we encountered challenges when reconstructing phylogenetic relationships among the different Bermin species. All 13 (9+4) species are clearly distinguishable in vivo in the lake, mainly by their distinct nuptial coloration, size, and body/mouth shape. However, we were unable to reconstruct their relationships using either the set of expressed transcripts, the set of BUSCO genes, or even whole mitogenomes retrieved from sequenced transcriptomic data. To provide an evolutionary context, we relied on relationships reconstructed using RAD-seq (multiple individuals per species, unpublished), which was the only method capable of distinguishing different species. This strongly resembles the situation observed in Victoria cichlids before the widespread application of whole-genome methods (Wagner et al., 2013).

We further compared the Bermin opsins within the context of other African cichlid lineages, focusing on key tuning amino acid sites known to shift the spectral (λmax) sensitivity of opsin pigments. Our results show that Bermin cichlids (and *Coptodon* species more broadly) consistently use the longer-wavelength sensitive amino acid variant for all opsins, except for SWS1 (the UV opsin), where both tuning sites retain the shorter-wavelength sensitive variant (Figure 1A). In other words, all cone opsins—except for one—exhibit a shift in sensitivity toward longer (red) wavelengths of light. Whether this pattern arises due to slightly higher turbidity, and thus a potentially red-shifted ambient light environment, or whether it reflects a phylogenetic constraint remains to be determined through a broader comparative analysis of other *Coptodons* and related genera.

### Opsin gene expression in Bermin cichlids and visual system in the deep-water species

Bermin cichlids have multichromatic colour vision, primarily utilizing five cone opsin genes: the short-wavelength-sensitive SWS2B and SWS2A found in single cones, and the middle-to-long-wavelength-sensitive RH2Aβ, RH2Aα, and LWS in double cones (Figure 2). The ratios of these opsins vary slightly, with one notable exception in the single cones of *C. flava* predominantly expressing the SWS2B gene, whereas all other species primarily express SWS2A (Figure 2). For comparison, the opsin gene repertoire of East African and Neotropical cichlid species typically includes three or four expressed cone opsin genes (Parry et al., 2005; Carleton et al., 2009; Escobar-Camacho et al., 2017). Overall, the opsin expression profile of Bermin cichlids is shifted toward longer wavelengths, a pattern commonly observed in cichlids inhabiting riverine or turbid lake environments (Halstenberg et al., 2005; Carleton et al., 2008; Hofmann et al., 2009a). Beyond cichlids, the dominance of LWS in double cones and SWS2A in single cones appears to be a widespread pattern among various teleosts and even non-teleost ray-finned fishes (Lupše et al., 2022; Musilova and Cortesi, 2023).

We then compared opsin expression in shallow- and deep-water Bermin species and found depth-associated differences in proportional opsin gene expression. While these differences are not as dramatic as in other systems—where deep-dwelling species exhibit complete opsin loss (e.g., Barombi Mbo; Musilova et al., 2019b)—we identified significant reductions in the expression of short-wavelength-sensitive opsins in the single cones of deep-water species (Table 2, Figure 3). Specifically, deep-water species express lower levels of SWS1 (UV-sensitive) and SWS2B (violet-sensitive), leading to an increased expression of SWS2A (blue-sensitive), likely reflecting the narrowed light spectrum at depth, where UV and violet wavelengths are largely absent. This trend aligns with patterns observed in other cichlid systems (e.g., Carruthers et al., in rev.) and mirrors the evolutionary loss of SWS1 and SWS2B in deep-sea fishes (Musilova et al., 2019a).

**Table 2.**
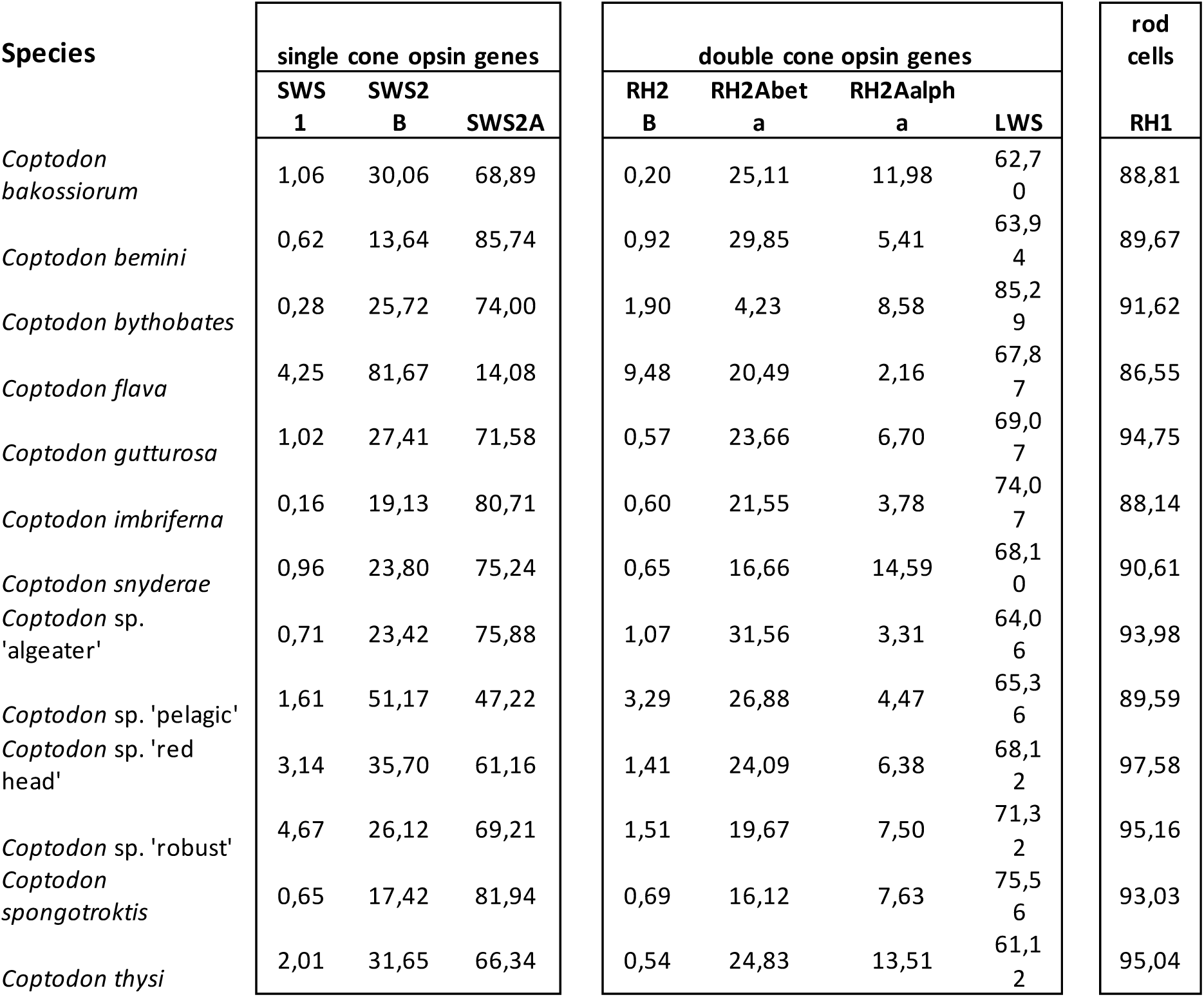
Average, proportional opsin gene expression in the single and double cones (in %).

**Figure 3:**
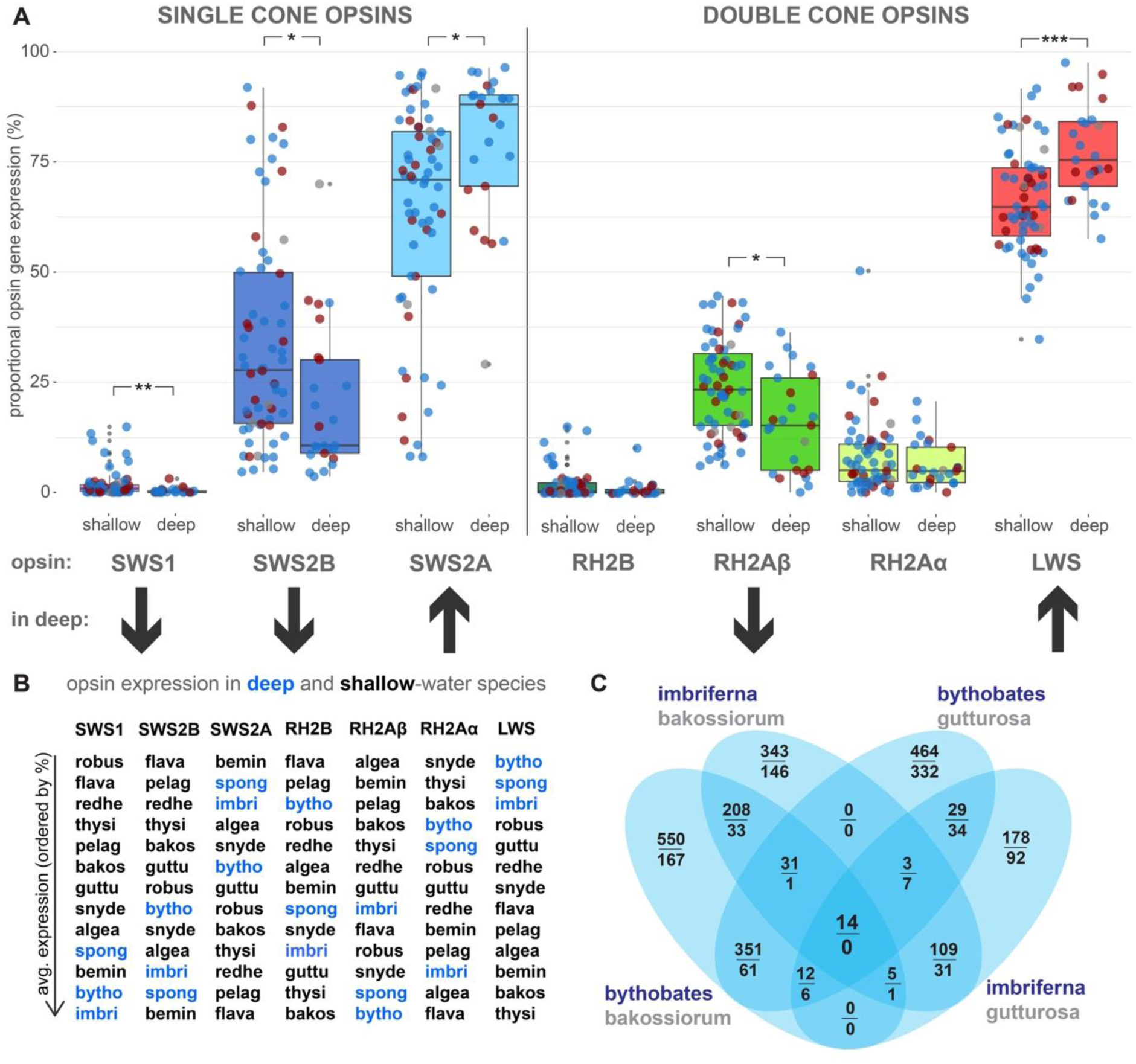
Opsin gene expression in shallow-water and deep-water species in the Bermin lake. A) Ten shallow-water species compared to the three species commonly or exclusively found in the deep. Visual opsin genes on x axis are ordered by their λmax sensitivity wavelength and the expression has been calculated separately for the single cone and the double cone cells in retina. We found lower expression of SWS1 (UV) and SWS2B (violet-sensitive) genes, which corresponds to the lack of shortest wavelengths in the deep, while the opposite pattern (increase, and not decrease as expected) has been found for the LWS (red) gene. B) Thirteen Bermin species ordered by average proportional expression for each visual opsin gene type. Blue highlights the deep-water species. Five opsins (SWS1, SWS2A, SWS2B, RH2Aβ and LWS) suggest association with the deep-water species and have been tested statistically (see Table 3, and Supplementary Table S5 and S6 for more details). C) Venn diagram showing the number of differentially expressed genes for the selected shallow-water and deep-water species pairs. See Table 4 for the list of differentially expressed genes.

Surprisingly, however, we observed the opposite trend at the long-wavelength end of the spectrum. The red-sensitive opsin LWS exhibited increased proportional expression in deep-water species (Figure 3, Table 2, Table 3, and Supplementary Tables S4, S5 and S6). In most other deep-water systems, LWS expression is reduced (e.g. in Masoko—Carruthers et al., in rev.) or even completely lost (e.g., Barombi Mbo—Musilova et al., 2019b; Tanganyika—Ricci et al., 2023). This reduction is typically attributed to the absence of red wavelengths at greater depths, as both ends of the light spectrum diminish with increasing water depth.

**Table 3.**
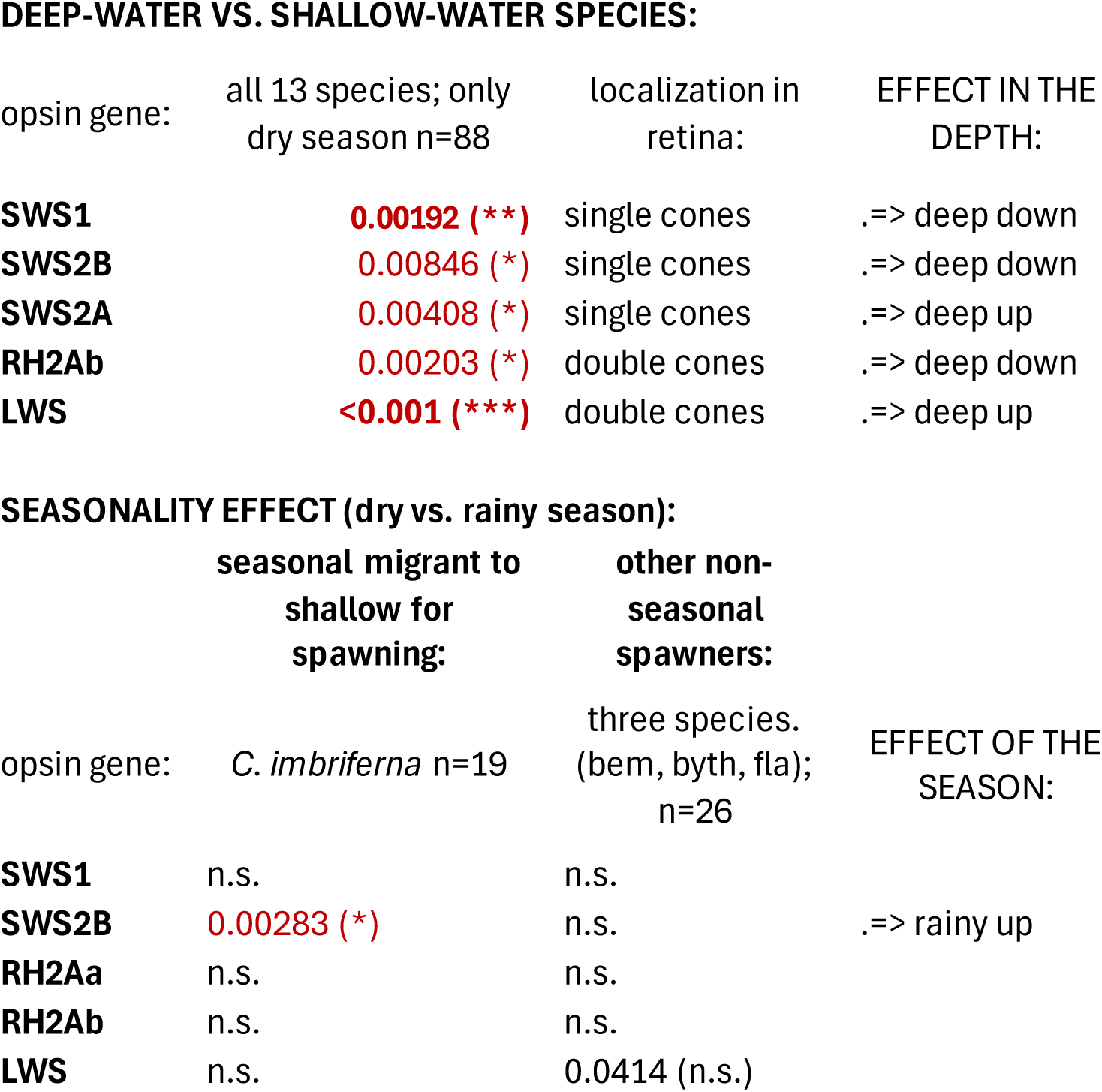
Formal statistical tests by beta-regression. P-values and level of significance after correction (in brackets).

**DEEP-WATER VS. SHALLOW-WATER SPECIES:**
| opsin gene: | all 13 species; only<br>dry season n=88 | localization in<br>retina: | EFFECT IN THE<br>DEPTH: |
| --- | --- | --- | --- |
| **SWS1** | **0.00192 (\*\*)** | single cones | .=> deep down |
| **SWS2B** | **0.00846 (\*)** | single cones | .=> deep down |
| **SWS2A** | **0.00408 (\*)** | single cones | .=> deep up |
| **RH2Ab** | **0.00203 (\*)** | double cones | .=> deep down |
| **LWS** | **<0.001 (\*\*\*)** | double cones | .=> deep up |

**SEASONALITY EFFECT (dry vs. rainy season):**
|  | seasonal migrant to<br>shallow for<br>spawning: | other non-seasonal<br>spawners: |  |
| --- | --- | --- | --- |
| opsin gene: | *C. imbriferna* n=19 | three species.<br>(bem, byth, fla);<br>n=26 | EFFECT OF THE<br>SEASON: |
| **SWS1** | n.s. | n.s. |  |
| **SWS2B** | **0.00283 (\*)** | n.s. | .=> rainy up |
| **RH2Aa** | n.s. | n.s. |  |
| **RH2Ab** | n.s. | n.s. |  |
| **LWS** | n.s. | 0.0414 (n.s.) |  |

The reason for the unexpected upregulation of LWS in Bermin deep-water species remains unclear. One possibility is that water clarity in Bermin is lower than in Barombi Mbo or Tanganyika, although no detailed optical measurements have been conducted in this lake. Additionally, the Bermin lake is shallower than Barombi Mbo (>110 m, with fish observed below 30 m), with a maximum depth of only 14 m (where all three deep-water species have been observed). As a result, the spectral filtering effects of the water column may not be as pronounced as in deeper systems (Hofmann et al., 2009; Carleton et al., 2009). Another potential explanation is the role of breeding colouration—many *Coptodon* species exhibit yellow and red hues, which may constrain selection against red sensitivity loss. Finally, this pattern may also stem from the unique evolutionary history of the Bermin species flock. All Bermin cichlids belong to the genus *Coptodon*, which is only distantly related to the Oreochromini lineage (i.e. the lineage with Barombi species) (Schedel et al., 2019; Astudillo-Clavijo et al., 2023). For these reasons, Bermin cichlids represent an important addition to comparative studies alongside oreochromine (Barombi), astatotilapiine (Masoko), and Neotropical heroine cichlids from Nicaragua.

Interestingly, the observed gene expression changes associated with the deep-water environment are more pronounced in males for single cone opsins (SWS1, SWS2A, SWS2B), whereas in double cone opsins (RH2s and LWS) the effect is primarily driven by females. This pattern could be a result of stochastic variation due to the lower sample size when analyzing a single sex, or it may indicate that the depth effect is influenced by additional factors in one or the other sex. For example, testosterone levels have been shown to affect the RH2-to-LWS ratio in lizard double cones (Tseng et al, 2018), and sexual maturity influences opsin expression in both male and female sticklebacks (Shao et al., 2014). While our sampling was not specifically controlled for reproductive state, all sampled individuals were confirmed to be adults.

The spectral sensitivity of visual pigments can also be altered by switching between retinal chromophores: a shift from A1 (11–cis retinal) to A2 (11-cis-3,4-dehydroretinal) moves the opsin λmax toward longer wavelengths. The use of A2 or a combination of both A1 and A2 has been documented in several freshwater fish, including cichlids (Carleton et al., 2008; Terai et al., 2017), and is also found in deep-sea fishes adapted to red-light sensitivity (Douglas et al., 2000). The cyp27c1 gene, a molecular marker for A2 synthesis, was identified in zebrafish (Enright et al., 2015) and can serve as a proxy for detecting A2 retinal in transcriptomic data. To determine whether A2 plays a role in *Coptodon* vision, we analyzed the expression of the cyp27c1 synthesis marker but found no evidence of its expression in the retina transcriptomes—regardless of sex or season. This strongly suggests that Bermin cichlids exclusively utilize A1 retinal in their visual pigments.

### Comparison of visual systems in Bermin and Barombi Mbo cichlid radiations

The two Cameroonian crater lakes, each hosting an endemic adaptive radiation, warrant a more detailed comparison (Figure 5). Both lakes are inhabited by a similar number of species— eleven in Barombi Mbo and thirteen in Bermin (nine valid + four undescribed)—as well as a comparable number of deep-water species. Both have one strict deep-water specialist (*Konia dikume* in Barombi, *Coptodon bythobates* in Bermin) and one seasonal migratory species (*Myaka myaka* in Barombi, *C. imbriferna* in Bermin). The phylogenetic origins of cichlids differ between the lakes (tribe Oreochromini in Barombi, Coptodonini in Bermin), as do their modes of reproduction (mouthbrooders in Barombi, substrate spawners in Bermin). Further differences include the striking yellow and sometimes red coloration of Bermin cichlids during breeding, whereas Barombi cichlids do not show much colour change during reproduction. The directions of adaptations in the visual system also differ: Barombi cichlids exhibit a larger divergence in double cones overall (mainly due to the loss of LWS cones in deep-water species), whereas Bermin cichlids show greater variability along the single cone axis (Figure 5). A particularly striking difference is the aforementioned adaptation of deep-water species: Barombi cichlids shift their double cones towards shorter wavelengths (due to the replacement of red-sensitive LWS with green-sensitive RH2Aα), whereas Bermin cichlids shift towards longer wavelengths in their double cones (due to increased LWS expression) (Figure 5).

**Figure 4.**
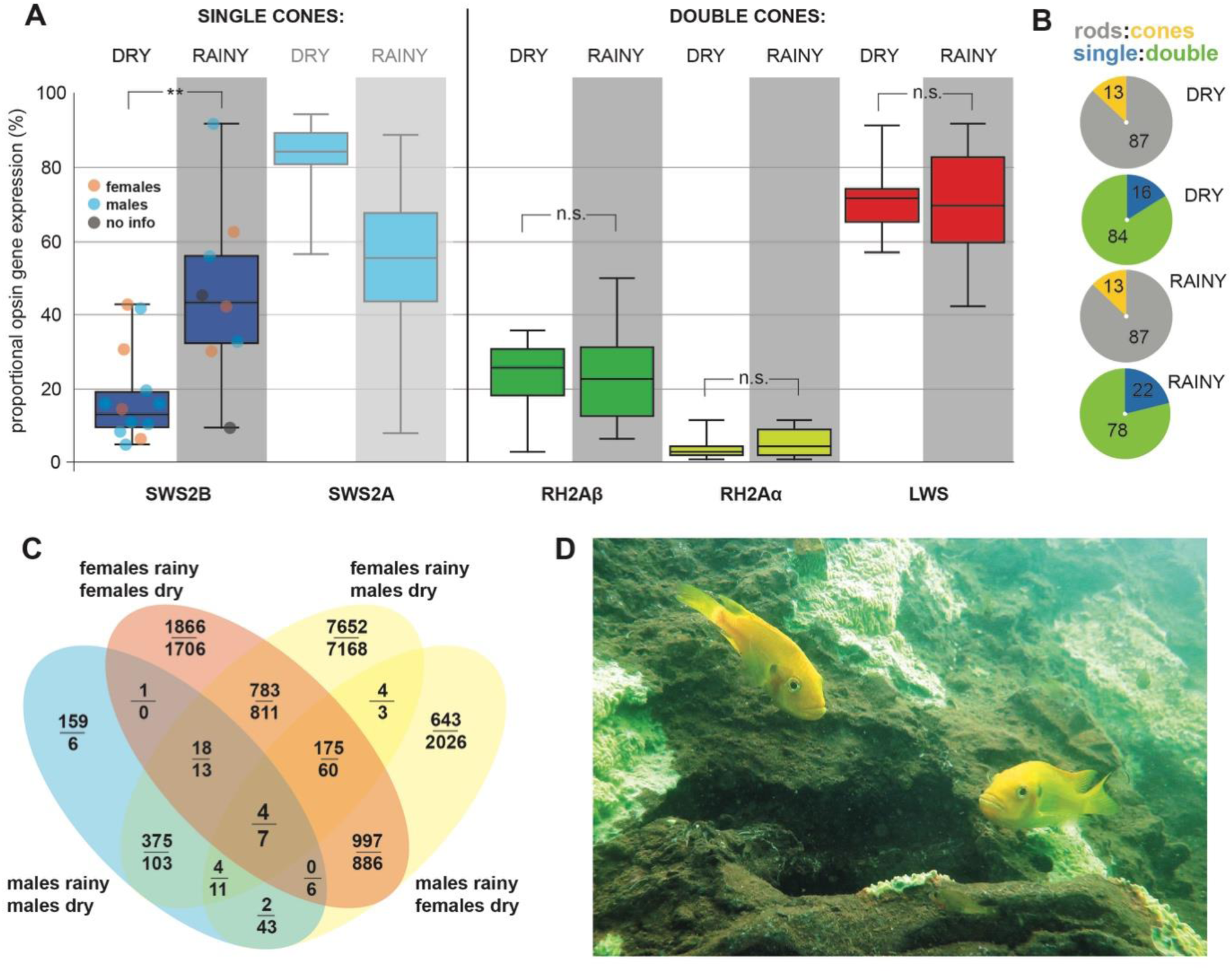
Differences in the visual system of *Coptodon imbriferna*, the species with seasonal spawning and depth preferences, between seasons. This species resides only in the deeper sections of the lake in the dry season (November to March) and migrates for spawning to the shallow water in the rainy season (May to August). A) The composition of visual opsins differs between the dry and rainy season in increased SWS2B to SWS2A ratio in single cones, a trend comparable to deep-water vs. shallow-water species. B) The rod:cone and single-cone:double-cone opsins in the dry and rainy season with no detected difference. C) Differential expression analysis on the retina transcriptomes revealed few candidate genes putatively associated with seasonal changes and migration for spawning period in *C. imbriferna* (for the list of genes see Table 4 and Supplementary Table S7). D) Breeding pair of *C. imbriferna* in the yellow spawning colouration.

**Figure 5.**
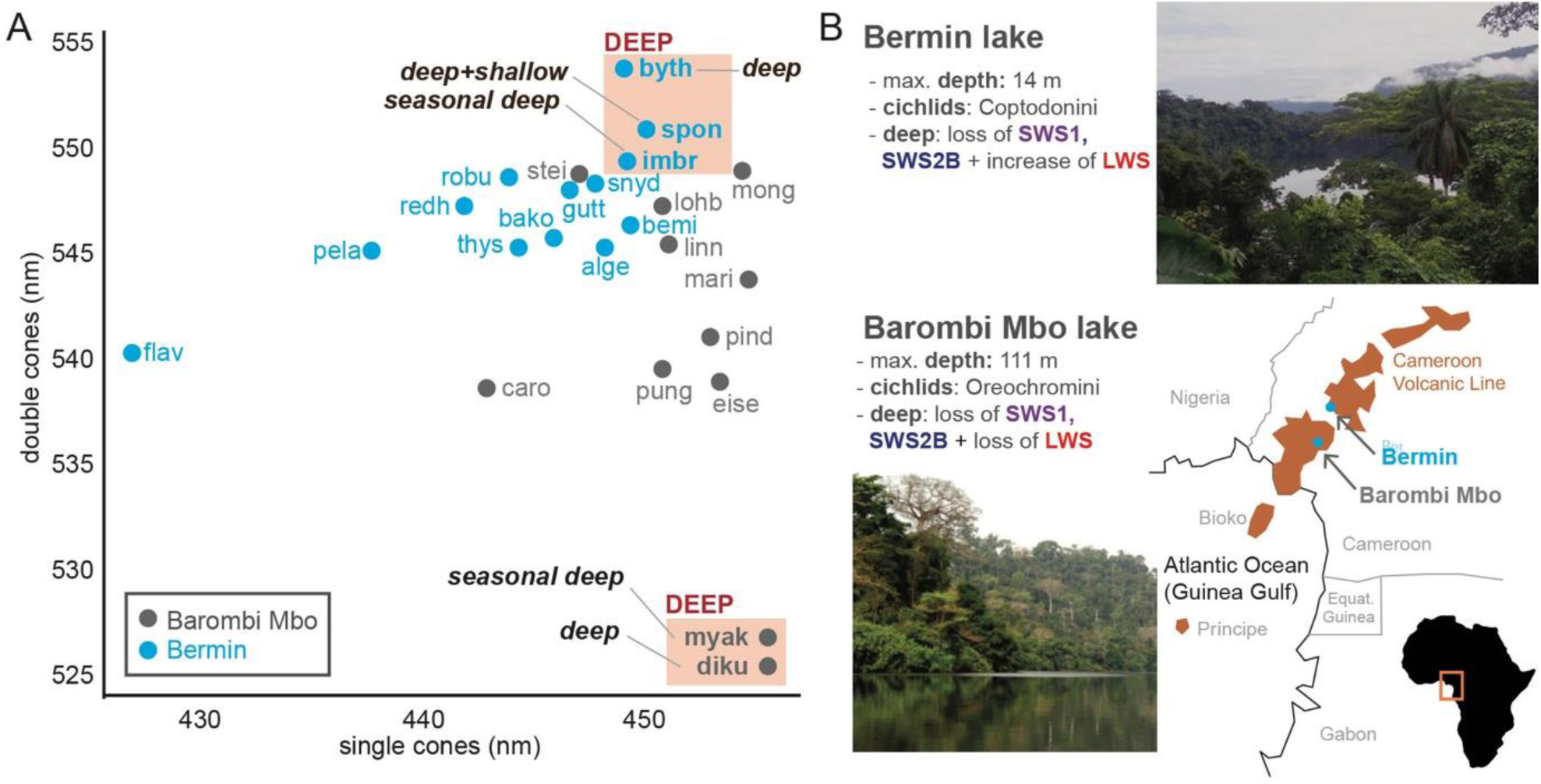
Overall retinal sensitivity and comparison between Bermin and Barombi Mbo cichlids. A) Note the opposite direction of adaptation in the deep-water species between the two cichlid systems. Double cones seem to shift towards shorter wavelenght in Barombi Mbo and longer wavelength in Bermin (compared to the rest of the species), a trend mainly driven by the expression of LWS opsin. Single-cone opsins, contrarily, tend to show the same trend of losing/lowering the expression of the shortest-sensitive opsins in both lakes, shifting the single cones towards longer wavelengths (= i.e., the center of the light spectrum). The overall retina sensitivity plot of the visual system are based on the weighted average of all seven opsin genes expressed within the single and the double cones separately, following the equation in Hofman et al.2009, and using experimentally measured values of cone opsin genes spectral sensitivity λmax from *Oreochromis niloticus* reference (Spady et al., 2006). Data for 11 Barombi species taken from Musilova et al. (2019b). B) Comparison of two crater lakes located in South-West Cameroon in West Africa. Barombi Mbo is a deeper lake hosting a radiation of oreochromine mouthbrooder cichlids, while Bermin is shallower and hosts endemic species flock of coptodonine substrate-spawning cichlids.

### Plasticity of the opsin gene expression in the seasonally spawning and migratory species

The seasonally deep-water species, *Coptodon imbriferna*, resides in the deep during the dry season, while it migrates to the shallow waters for spawning in the rainy season (Stiassny, 1992). The expression profiles show that the visual system differs between seasons in the ratio of SWS2B to SWS2A expression in the single cones (Table 3, Figure 4), while no changes are observed for the other genes. The increased SWS2A:SWS2B ratio in the single cones corresponds to the same direction as in the comparison of the deep- and shallow-water species and aligns with other cichlid studies focusing on depth (Carruthers et al., in rev.; Musilova et al. 2019b). The seasonal migration could be seen as a functional parallel to the deep vs. shallow-water species comparison, and the direction of change is the same for the SWS2B gene. Hence, the observed plasticity of the visual system in the seasonal migratory species may also represent a mechanism by which the visual system adapts: a plastic response to the environment, which may subsequently lead to fixed differences among species. We further tested whether seasonal differences could be found in other non-seasonal species in the lake, and no differences were observed for three other species sampled in both seasons (Supplementary Table S6). Therefore, the seasonal differences are found only in a species with seasonal reproduction (*C. imbriferna*) and the related change in depth. Interestingly, when *C. imbriferna* samples are analyzed separately by sex, mild differences in the expression of RH2Aα (males) or RH2Aβ (females) are also observed between seasons (Supplementary Table S6); however, the sample size was unfortunately rather low for this test. Similarly, to check for differences between sexes, we complemented our seasonality analysis in *C. imbriferna* by testing the effect of sex in the dry season samples of this species. We found differences in the RH2A:LWS ratios (Supplementary Table S6). No differences were found when testing samples from the rainy season, as well as in four other species with male and female samples (Supplementary Table S6). However, the samples were not specifically collected for this test, and hence the sample size was limited, reducing the power of these two analyses.

### Differential gene expression of retinal transcriptomes: genes associated with deep-water species, and seasonally relevant

We took advantage of the sequenced retina transcriptomes and searched for other putative candidate genes associated with adaptation to the deep-water environment, as well as with seasonality in *C. imbriferna*.

To search for the deep-water associated candidates, we used the retina transcriptomes of two deep-water species (*C. bythobates* and *C. imbriferna*) and two selected shallow-water species (*C. bakossiorum* and *C. gutturosa*). We were specifically interested in genes that were differentially expressed in all pairwise (i.e., shallow-deep) comparisons, as this design should reveal the putative candidate genes associated with depth. We identified 14 differentially expressed genes, all of which were upregulated in the deep-water species (Figure 3C, Table 4, Supplementary Table S7). Interestingly, in a similar comparison in the Masoko cichlids, all identified differentially expressed genes were also upregulated (i.e. no downregulated genes were found) in the deep-water ecomorph (Carruthers et al., in rev.). Of the 14 identified differentially expressed genes in Bermin cichlids, five are putatively associated with vision and are involved in retinal repair and differentiation, cone development, or blood flow control (Table 3; Supplementary Table S7). Specifically, HEY1 (hes-related bHLH transcription factor) is involved in regulating the regenerative response of Müller glia in the retina, as tested in zebrafish after injury (Sahu et al., 2021; Maier & Gessler 2000). Endothelin-1 (ET-1) is involved in blood flow control in the retina (Salvatore et al., 2010; Freeman et al., 2014) and photoreceptor synaptic transmission (Delyfer et al., 2005). Fibrillin 1 (fbn1) is a crucial ocular glycoprotein and plays a role in the structural and functional stability of retinal arterioles (Hubmacher et al., 2014; Alonso et al., 2023). And collagen col25a1 is associated with the structure and function of direction-selective retinal ganglion cells (Deng et al., 2016; Sangsin et al., 2016), which are essential for motion detection and orientation (Barlow et al., 1963; Vaney et al., 2012; Baden et al., 2016; Rheaume et al., 2018; Jiang et al., 2022) (Table 3 and Supplementary Table S7). Interestingly, none of the differentially expressed genes found between shallow and deep-water species overlaps either with the genes found in the Barombi Mbo analysis using the same design (Musilova et al., 2019b), nor with the DE analysis in the deep- and shallow-water Masoko ecomorphs (Carruthers et al., in rev.), although the functional categories do overlap. This suggests that major molecular mechanisms may not be conserved across different cichlid systems, in contrast to smaller-scale mechanisms, such as opsin expression tuning. The DGE analysis across systems is also somewhat limited due to the varying methods and teams involved in sample collection, and likely due to other factors that may obscure the signal.

**Table 4.**
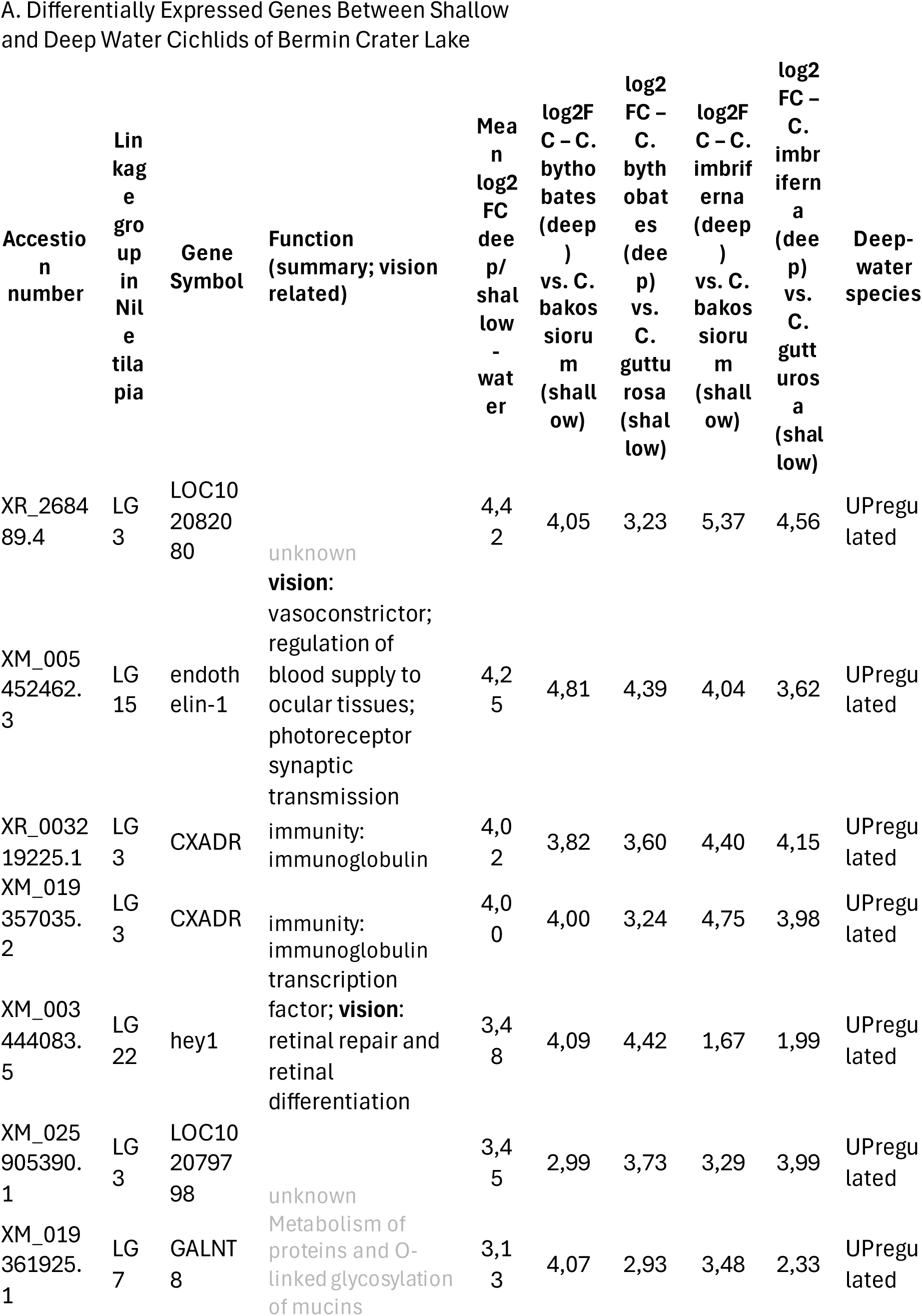
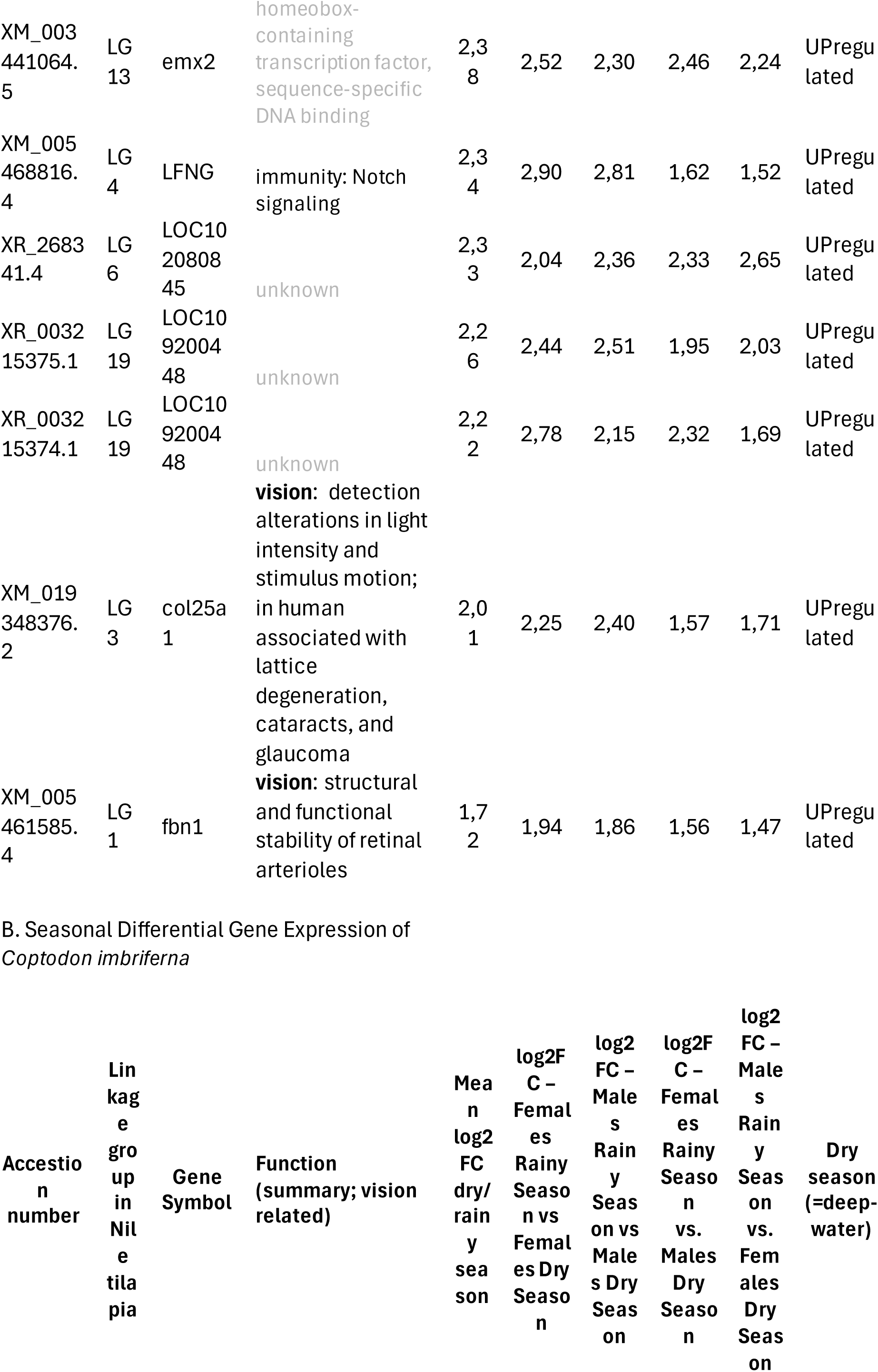
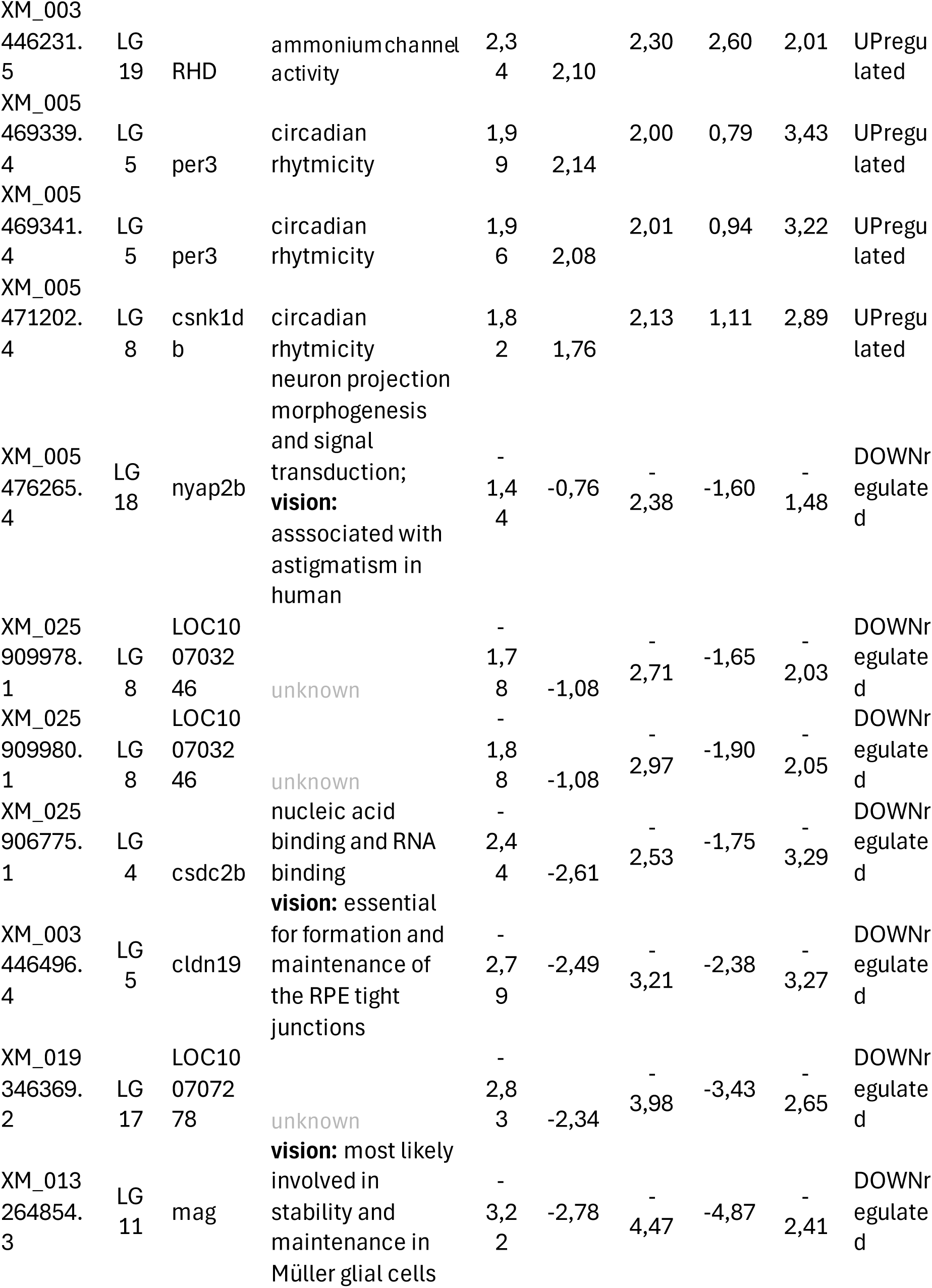
Differential Gene Expression Analysis in Cichlids of Bermin Crater Lake.

Focusing on seasonality, we analyzed the dry and rainy season transcriptomes of *C. imbriferna*, with sexes analyzed separately (Figure 4C, Table 4, Supplementary Table S7). We aimed to similarly compare all pairwise comparisons (i.e., dry vs. rainy), and identified eleven genes that were differentially expressed across comparisons (four upregulated and seven downregulated during the dry (=deep-water) season). Among these, two downregulated genes are associated with visual processes: MAG (myelin-associated glycoprotein), which is involved in neural regeneration and structural stabilization via Müller glial cell maintenance (Reichenbach and Bringmann 2010; Bringmann et al., 2009); and claudin-19 (CLDN19), which is associated with tight junctions in the blood-retinal barrier and is critical for retinal pigment epithelium (RPE) integrity and retinal homeostasis (Naylor et al., 2019). Two upregulated genes, PER3 and CSNK1Db, are known to be related to circadian rhythms (Constance et al., 2005). Further, we tested whether any of the differentially expressed genes overlapped with those found in the deep-vs.-shallow analysis. Comparing the two seasons of *C. imbriferna* essentially means comparing when the species inhabits the deep and shallow environments. Surprisingly, we did not detect any overlap between the two analyses suggesting that the similar molecular trajectory is not used in the plastic response to the depth (seasonal species) and the long-life adaptation (deep-water species). However, we did find similarities between the seasonality analysis and the deep-vs-shallow analysis in Barombi Mbo, namely the upregulation of several circadian rhythm genes (although not the same ones) in the Barombi deep-water species and in *C. imbriferna* during the dry season (=deep). Circadian genes are known to change expression throughout the day (e.g., Bolton et al., 2021), and the two genes reported as upregulated in the dry (=deep) season, per3 and csnk1db, are both known to have a low peak of expression during the daytime. This possibly suggests less prominent periodicity in expression, or compensation for the lack of light in the deep habitat. Although we haven’t specifically controlled for the time of day during sample collection, both shallow- and deep-water samples were typically collected around the same time of day, irrespective of the season. The expression of circadian genes in the deep-water habitat therefore deserves specific focus in future studies. Circadian genes commonly appear among the differentially regulated genes in deep-vs-shallow water comparisons, alongside other categories such as immunity genes, hypoxia tolerance, and maintenance and endurance of visual system tissues (Musilova et al., 2019b; Ricci et al., 2023).

## Conclusion

In this study, we provide a comprehensive analysis of visual system evolution across the entire adaptive radiation of Bermin crater lake cichlids. By including all 13 species (nine described and four undescribed), we identify a clear association between opsin expression profiles and habitat depth. Additionally, we examine seasonal variations in the visual system of a species that inhabits deep water during the dry season and shallow water during the rainy season. We identify both shared and unique adaptations to deep-water environments, offering insights into the broader mechanisms of cichlid visual evolution. A typical deep-water adaptation is observed in the single cones of the retina, characterized by an increased expression ratio of short-wavelength-sensitive opsins (SWS2A:SWS2B). However, we also detect an unexpected increase in the expression of the red-sensitive LWS opsin, suggesting a unique visual adaptation. Beyond opsins, we report an upregulation of circadian rhythm genes in the seasonally deep-water species during the dry season, when it inhabits deeper waters, compared to the rainy season, when it occupies shallow habitats.

## Supporting information

S1 - Opsin gene expression profiles: Opsin gene expression profiles based on the retina transcriptomes of 13 cichlid species from Bermin crater lake.

Supplementary_tables_Klodawska et al

## Acknowledgements

We extend our deepest gratitude to the Bakossi people for their warm hospitality, guidance, and permission to conduct our research on their land. We specifically want to thank to Ewange Marcus Alung, Esong Frankline Ejome, his son Ejome Ekolle Clovis, and their families for their invaluable support and friendship throughout our study. We further thank Antoine Pariselle and Ngando Ephesians Jiku for fieldwork assistance, and Cyrille Dening for logistical support. We would also like to express our appreciation to our colleagues Gina Sommer, Veronika Truhlářová, Samuel Didier Njom, and Gabriela Wofková for their unwavering support, inspiring conversations, and companionship during our fieldwork, lab work, and beyond.

## Supplementary Material

**Figure S1 -** Opsin gene expression profiles: Opsin gene expression profiles based on the retina transcriptomes of 13 cichlid species from Bermin crater lake. Box plots showing the median and interquartile range (IQR) of 25% to 75% with outliers outside the 1.5 IQR are based on 5 – 11 individuals per species. See Table T1 for more details.

**Table S1 -** Amino acid sites in the opsin genes across African cichlids

**Table S2 -** Samples used for the Sanger sequencing

**Table S3 -** Primers used for the Sanger sequencing of the single opsin genes

**Table S4 -** Raw opsin gene expression data for all individuals

**Table S5 -** Opsin gene expression per gene and per species with the test for the deep-water species

**Table S6 -** Formal statistical beta-regression tests

**Table S7 -** Differential Gene Expression Analysis in Cichlids of Bermin Crater Lake - detailed version

