## Supplementary material for "Seeing in the deep: evolution of the opsin gene expression in Bermin crater lake cichlids": S1 - Opsin gene expression profiles: Opsin gene expression profiles based on the retina transcriptomes of 13 cichlid species from Bermin crater lake.

*Coptodon* sp. "algeater":

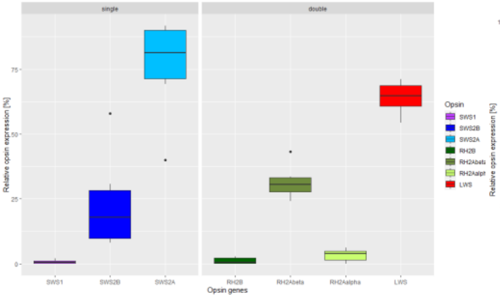


*Coptodon bakossiorum*:

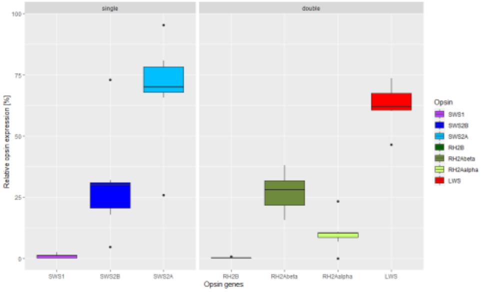


*Coptodon bemini*:

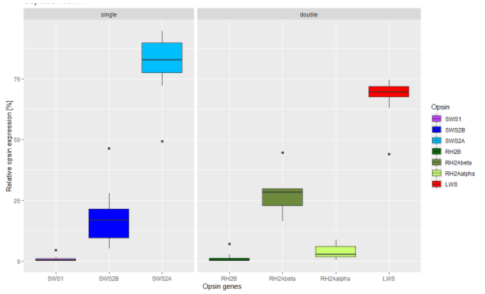


*Coptodon bythobates*:

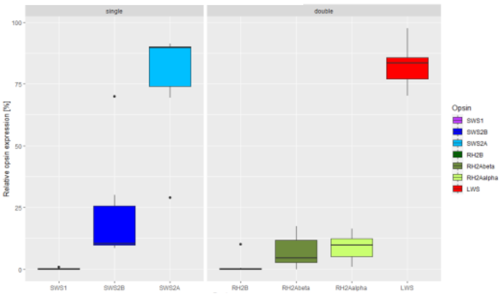


*Coptodon flava*:

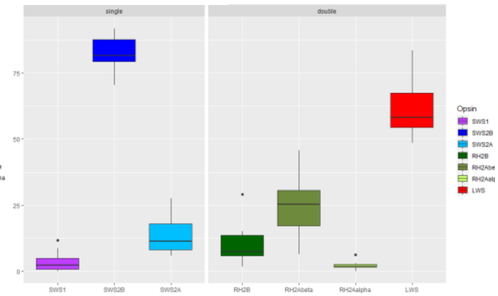


*Coptodon gutturosa*:

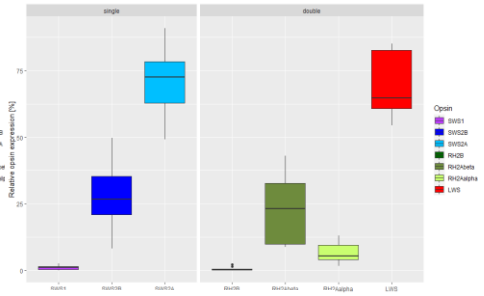


*Coptodon imbriferna*:

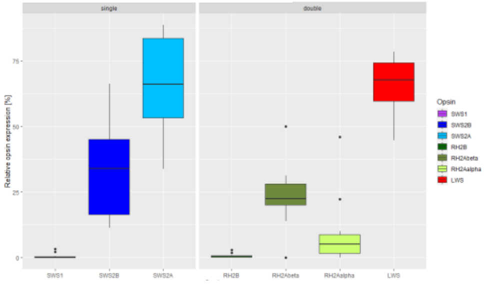


*Coptodon* sp. "pelagic":

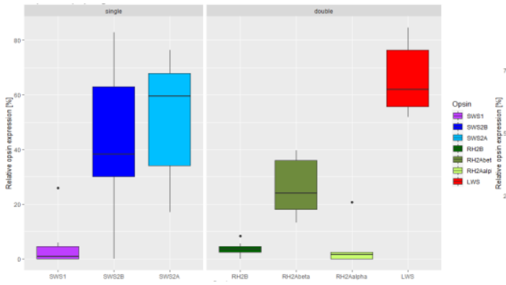


*Coptodon* sp. "redhead":

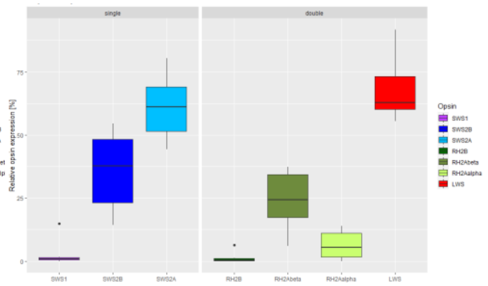


*Coptodon* sp. "robust":

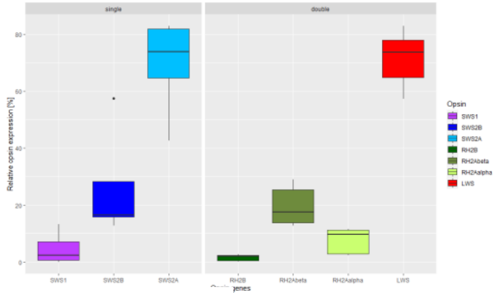


*Coptodon snyderae*:

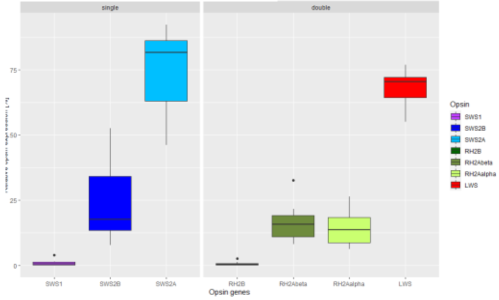


*Coptodon spongotroktis*:

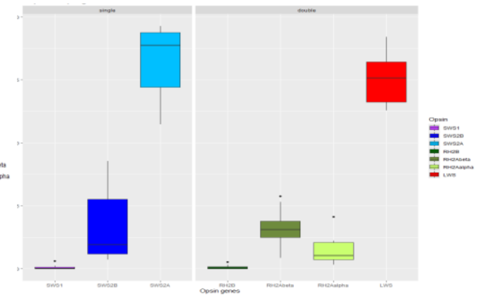


*Coptodon thysi*:

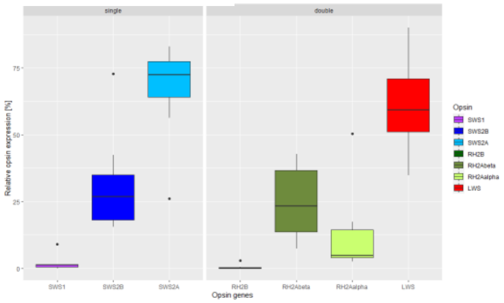
